# Methionine dependence in cancer cells due to lack of B_12_-dependent methionine synthase activity

**DOI:** 10.1101/2025.02.27.640554

**Authors:** Mohamed M.A. El Husseiny, Roland Nilsson

**Author notes:** Corresponding author: Roland Nilsson.

## Abstract

Human cells can synthesize methionine from homocysteine and folate-coupled methyl groups via the B_12_-dependent enzyme methionine synthase (MTR). Yet, it has been known for decades that cancer cells fail to grow when methionine is replaced by homocysteine, a phenomenon known as methionine dependence. The underlying mechanism remains unknown. Here, we report evidence that methionine dependence is caused by low MTR activity secondary to a B_12_ deficiency. High levels of the B_12_ cofactor were required to revert methionine-dependent cancer cells to grow on homocysteine. The adapted “revertant” cells display gene expression signatures consistent with reduced invasion and metastasis. Metabolic flux analysis indicated that methionine-dependent cells do not fully activate MTR when cultured in homocysteine. High concentrations of homocysteine partially rescued growth of methionine-dependent cells. Expression of a B_12_-independent methionine synthase enzyme in cancer cells restored growth on homocysteine and normalized the SAM:SAH ratio, while overexpression of the B_12_-dependent human enzyme had no effect. These data establish a biochemical basis for the long-standing question of methionine dependence in cancer, and may open up new avenues for exploiting the phenomenon for cancer therapy.

## Introduction

Methionine plays a central role in human cellular metabolism as a proteinogenic amino acid, a major source of methyl groups and polyamines, and a precursor of cysteine (Fig. 1a). The “activated” form of methionine, S-adenosyl-methionine (SAM), is the central methyl donor for methylation of DNA, RNA, proteins, lipids and other small molecules. Methionine can be synthesized from homocysteine and a methyl group carried by tetrahydrofolate (CH_3_-THF), and certain cell types, in particular fibroblasts, thrive on medium where methionine is replaced by homocysteine, demonstrating that homocysteine is sufficient for their biosynthetic needs. Yet, it has been observed for decades that a variety of cancer cell lines fail to proliferate in homocysteine medium ^1,2^, a phenomenon termed “methionine dependence”. In animal models, reducing the plasma methionine level through a methionine-restricted diet has repeatedly been shown to reduce growth of primary tumors and suppress metastasis ^3–8^, indicating that methionine dependence occurs also *in vivo*.

**Figure 1.**
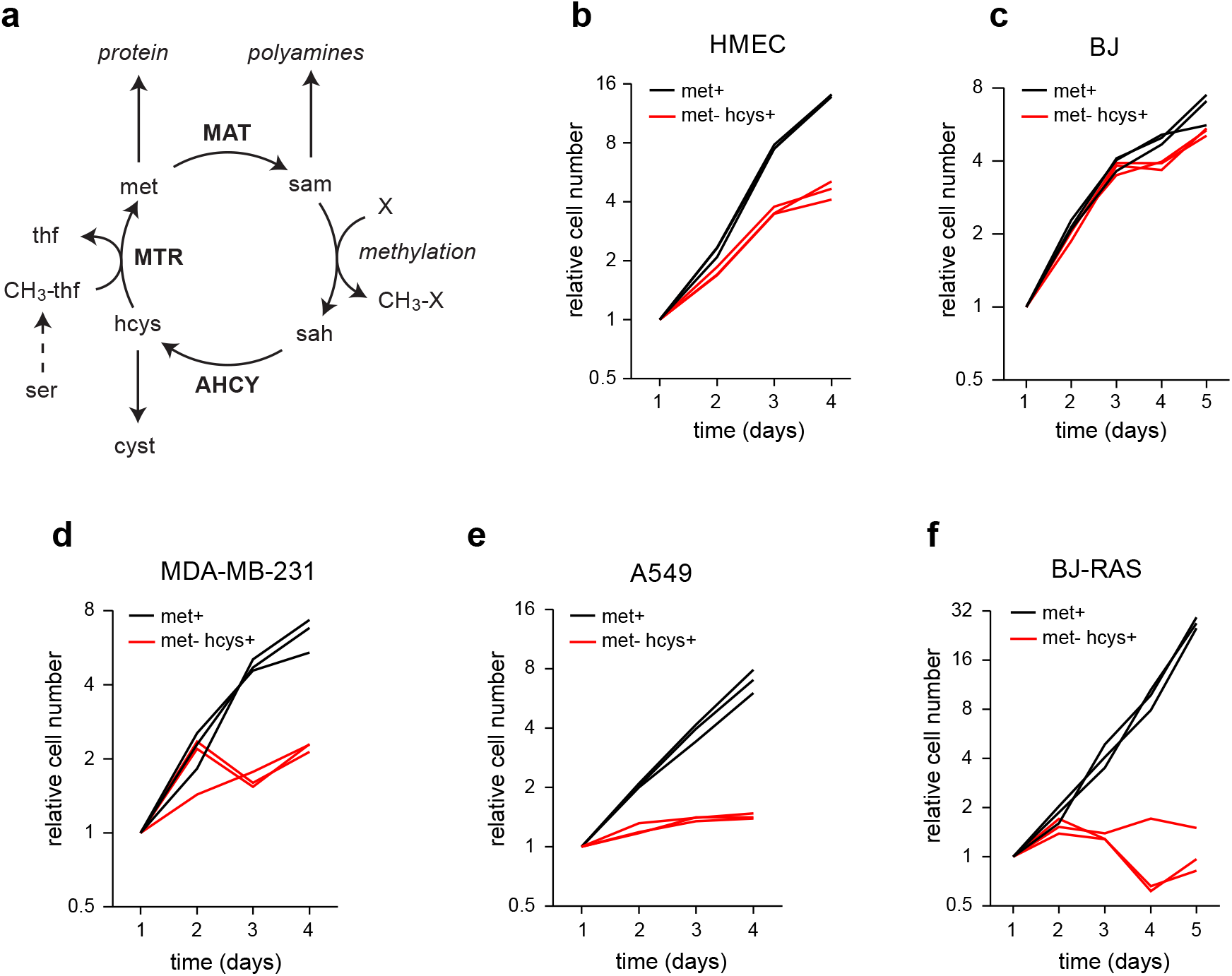
Methionine dependence occurs in tumor-derived cancer cells and oncogene-transformed cells. **a**, Schematic representation of methionine metabolism. AHCY, adenosylhomocysteinas; MAT, methionine adenosyltransferase; MTR, methionine synthase; met, methionine; sam, S-adenosylmethioine; sah, S-adenosylhomocystine; hcys, homocysteine; cyst, cystathionine; thf, tetrahydrofolate; CH_3_-thf, 5-methyl-tetrahydrofolate. **b–e**, Growth curves for normal human mammary epithelial cells (HMEC), human foreskin fibroblasts (BJ), breast cancer cells (MDA-MB-231), lung cancer cells (A549), and BJ cells transformed with the SV40 Large-T antigen and oncogenic HRAS^V12^ (BJ-RAS), in methionine-containing (met^+^) medium and methionine-free, homocysteine-containing medium (met^−^hcys^+^). Cell numbers relative to day 1 from three independent time course experiments are shown (n = 3).

Methionine dependence is not unique to tumor-derived cancer cells, but can also be induced in fibroblasts by transformation with the SV40 virus-derived large-T protein ^9^, and in mammary epithelial cells by expression of an oncogenic PI3-kinase ^10^, indicating that the mechanisms that cause cancer transformation also reprogram methionine metabolism. Hence, methionine dependence is not due to a genetic defect in cancer cells, such as loss of a critical enzyme during somatic evolution. Interestingly, methionine dependence also occurs naturally during embryonic development, where methionine deprivation of embryos prevents neural tube closure ^11^. Human embryonic stem cells also appear to be slightly methionine-dependent with limited growth on homocysteine ^12^. This suggests that methionine dependence is closely associated with a particular cellular differentiation state that occurs naturally in early development and re-appears in transformed cells.

While methionine dependence has been studied for decades, the underlying mechanism remains unclear. The obvious candidate would be a defect in methionine synthase (5-methyltetrahydrofolate-homocysteine methyltransferase, MTR; Fig 1a), but this hypothesis has been rejected by many investigators due to early reports that methionine-dependent cells have intact MTR activity ^13,14^. However, these studies only assayed MTR activity in cell lysates, and did not quantify metabolic flux through MTR in live cells. To our knowledge, the only quantitative data on MTR flux is from one study of fibrosarcoma cells in methionine medium using ^13^C isotope tracing ^15^. This study found low MTR activity, such that most homocysteine produced was released into the medium. In homocysteine medium, one study performed isotope tracing with deuterated homocysteine in breast cancer cells, but did not quantify metabolic fluxes ^16^. Thus, there is hardly any information on the metabolic response to homocysteine medium in either methionine-dependent or -independent cells. In this study, we therefore set out to characterize methionine metabolism in cancer cells and reassess possible metabolic mechanisms of methionine dependence.

## Results

### Methionine dependence in tumor-derived and oncogene-transformed cells

We began by characterizing the growth of various normal and transformed cells in methionine-free homocysteine (met^−^hcys^+^) medium compared to methionine-containing (met^+^) medium. Normal skin fibroblasts (BJ cells) proliferated in met^−^hcys^+^ medium at rates similar to that in met^+^ medium (Fig 1c), indicating that these cells can synthesize sufficient amounts of methionine from homocysteine. Normal mammary epithelial cells also proliferated in met^−^hcys^+^ medium, although somewhat slower than in met^+^ (Fig 1b). In contrast, a variety of cancer cell lines originating from breast, lung, brain and colon cancers failed to grow in met^−^hcys^+^ medium (Fig 1d,e, Supplementary Fig 1a–c), demonstrating that these cells are methionine-dependent. Some, but not all, cell lines exhibited growth during the first day in met^−^hcys^+^ medium before proliferation ceased (Fig 1d, Supplementary Fig 1c), possibly reflecting differences in cellular amino acid stores. Lack of growth in met^−^hcys^+^ was not due to homocysteine toxicity, since proliferation was unaffected in met^+^hcys^+^ medium (Suppl Fig 1d–f). We also observed that methionine dependence appeared in isogenic BJ cells transformed by expression of the SV40 Large-T antigen and oncogenic HRAS^V12^ (BJ-RAS) ^17^ (Fig 1f), confirming that MD can be induced by oncogenic signaling, and does not require loss of genetic elements. Taken together, this data supports the notion that methionine dependence is associated with cancer transformation across several cancer types.

### Reversion to methionine independence depends on vitamin B12

A fundamental problem in cancer therapeutics is the ability of cancer cells to evade treatments by activating alternative signaling or metabolic pathways in response to drugs. To investigate whether cancer cells are able to adapt to methionine deprivation, we performed long-term cultures in met^−^hcys^+^ medium. Remarkably, we observed no growth for up to four weeks in the tumor-derived cell lines tested, suggesting that cancer cells cannot easily bypass the mechanism that causes methionine dependence (Fig. 2a–c). The *in vitro* transformed BJ-RAS fibroblasts were an exception (Supplementary Fig. 2a), perhaps reflecting cell lineage differences. These results appeared to contradict previous reports of “revertant” cell lines generated through similar long-term methionine deprivation experiments ^9,18–20^. A systematic literature review revealed that almost all of these experiments used medium containing very high levels of vitamin B_12_ (cobalamin), a necessary cofactor for methionine synthase (Supplementary Table 1). We therefore performed long-term cultures with high levels of B_12_ and found that this strongly promoted reversion of MDA-MB-231 and MCF7 cells (Fig. 2a,b). In A549 cells, B_12_ supplementation did not promote full recovery (Fig 2c), but allowed cell colonies to form (Fig 2d, Supplementary Fig. 2c) that could later be expanded. These results suggest that at least in some cell types, methionine dependence is related to insufficiency of B_12_ for a process that is specifically required in met^−^hcys^+^ medium, such as methionine synthesis. Although high B_12_ allowed cells to survive in met^−^hcys^+^ medium, the resulting “revertant” cells still grew very slowly in met^−^hcys^+^ compared to met^+^ medium (Fig. 2e,f, Supplementary Fig. 2b), in agreement with previous reports (Supplementary Table 2). Thus, high exogenous B_12_ cannot fully overcome the underlying metabolic defect.

**Figure 2.**
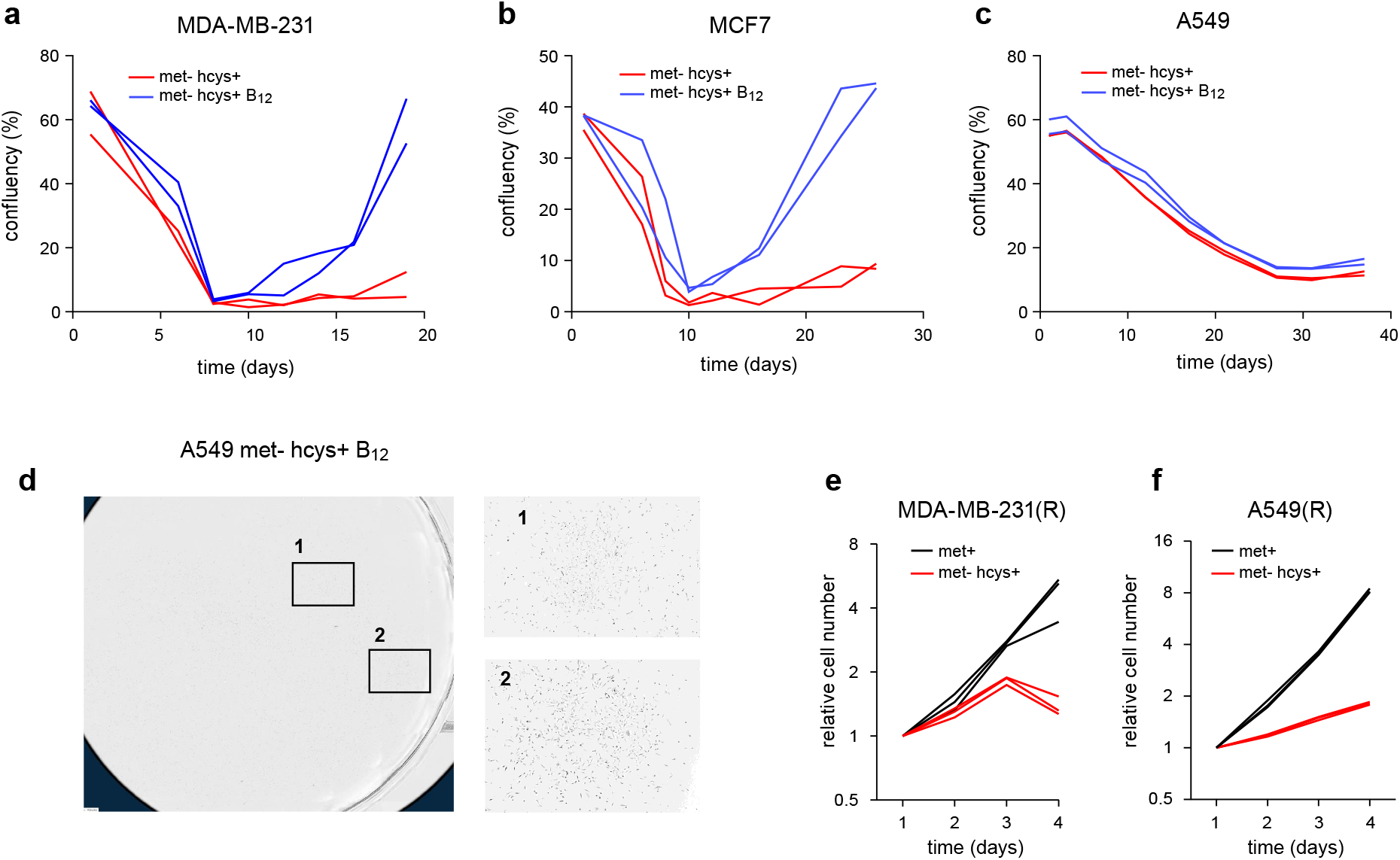
Reversion of methionine dependence in cancer cells requires vitamin B_12_. **a–c**, Long-term growth curves of breast cancer cells (MDA-MB-231, MCF7) and lung cancer cells (A549) in homocysteine-containing medium with either 0.003 µM (met^−^hcys^+^) or 1.5 µM vitamin B_12_ (met^−^ hcys^+^ B_12_). Confluency (%) from two independent cultures are shown (n=2). **c**, Example of cell colonies observed in A549 cells in met^−^hcys^+^ B_12_ medium. **e–f**, Growth curves of revertant cells MDA-MB-231(R) and A549(R) obtained from long term cultures, in methionine-containing (met^+^) medium and methionine-free, homocysteine-containing medium (met^−^hcys+). Cell numbers relative to day 1 from three independent time course experiments are shown (n = 3).

### Gene expression signatures of methionine dependence

To understand how reversion of methionine-dependent cells in the presence of B_12_ affects cell differentiation and gene expression programs, we next performed single-cell RNA-sequencing. We selected the MDA-MB-231 cell line, a commonly used model of triple-negative breast cancer ^21^, and analyzed single cell transcriptomes of parental cells before selection (D0) and “revertant” cells selected in met^−^hcys^+^ medium for 21 days (D21).

In D0 cells, single-cell expression analysis revealed two major clusters of cells (Fig. 3a). Cluster 0 exhibited gene expression signatures of cell differentiation and migration (Fig. 3b), while cluster 1 expressed extracellular matrix, wound healing and antigen presentation signatures (Fig. 3c), features reminiscent of fibroblasts or related mesenchymal cells. Further analysis of genes overexpressed in these clusters against a database of 1,355 human cell type-specific expression patterns ^22^ indicated that cluster 0 cells are most similar to endothelial cells, while cluster 1 cells are similar to mesenchymal cells (Supplementary Fig. 3c). A small third cluster consisted of cells expressing interferon-response genes (Fig. 3a, Supplementary Fig. 3a), which have previously been observed in cancer cells and associated with resistance to DNA damage ^23^; we excluded this cluster from further analyses. In D21 cells, cluster 0 decreased while cluster 1 became more prominent (Fig. 3d), suggesting that reversion to methionine independence promotes the fibroblast-like phenotype. Interestingly, genes differentially expressed between cluster 0 and cluster 1 closely matched a previously established signature ^24^ of a lung-metastasizing subpopulation MDA-MB-231 cells (Fig. 3e, Supplementary Fig 3b), suggesting that cluster 0 corresponds to the metastasis-capable subpopulation. Hence, reversion to methionine-independence appears to select against the more aggressive phenotype in MDA-MB-231 cells. These results are in line with reports that reversion is accompanied increased anchorage dependence ^16,18^, and underscores the close association between methionine dependence and cancer transformation.

**Figure 3.**
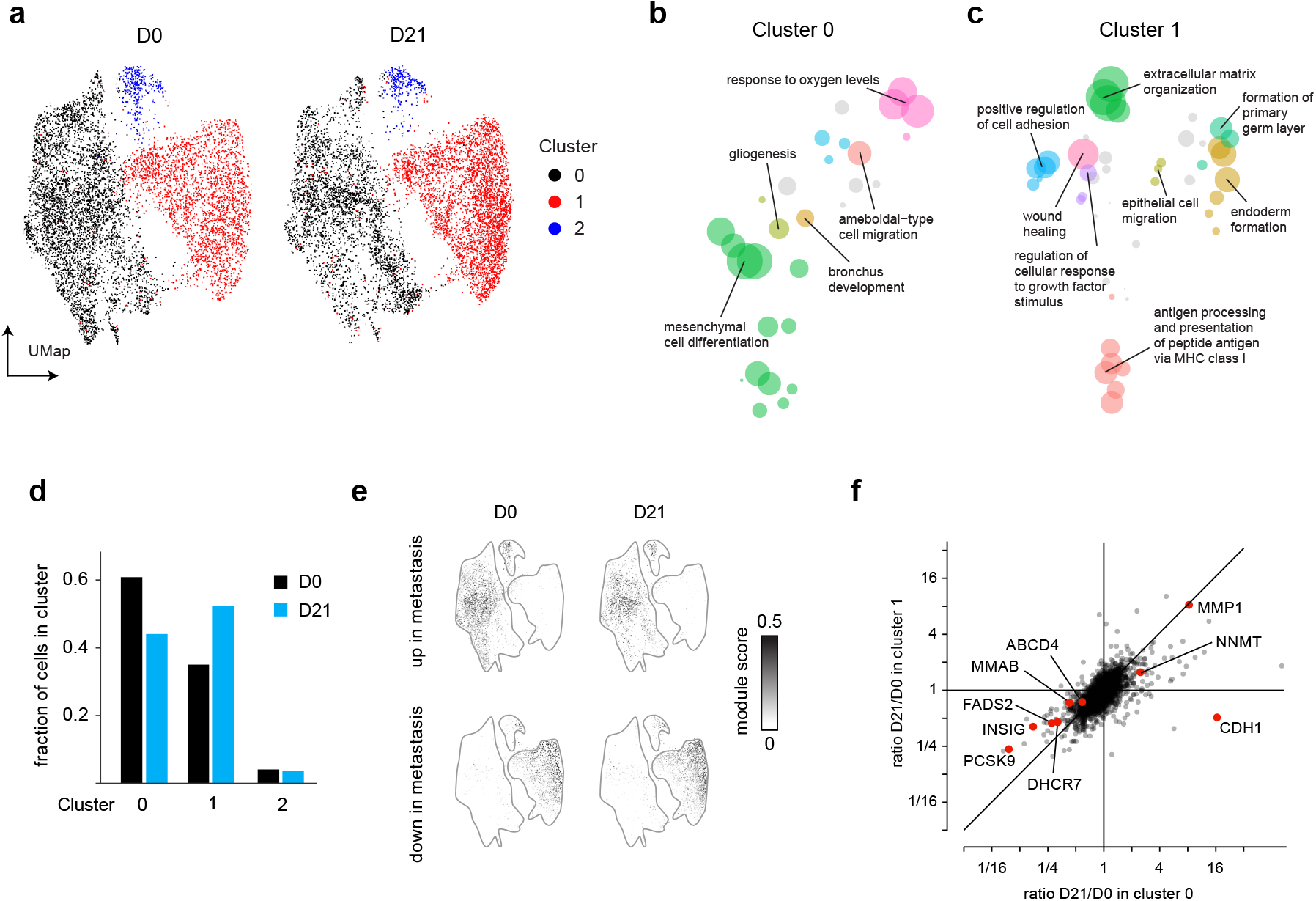
Cell differentiation and gene expression programs during reversion. **a**, UMap projection and clustering of MDA-MB-231 parental cells at day 0 (D0) and revertant cells at day 21 (D21). **b–c**, Non-redundant gene ontology (GO) pathways enriched in cluster 0 and 1, visualized as UMap projection of semantic similarity. Size of circles represent pathway over-representation score. **d**, Fraction of cells belonging to each cluster in D0 and D21 cells. **e**, Association between cell clusters and a previously described metastatis gene signature, shown as module score of genes up-regulated (top) and down-regulated in metastasis (bottom) for each cell. **f**, Scatter plot of gene expression fold change between D0 and D21, in cluster 0 and 1.

When comparing revertant cells to parental, the most prominent change was decreased expression in both clusters 0 and 1 of genes in sterol synthesis, fatty acid synthesis and lipoprotein trafficking, such as DHCR7, FADS2 and PCKS9 (Fig. 3f). Gene set enrichment analysis confirmed downregulation of these pathways (Supplementary Table 3). These may be direct effects of methionine insufficiency, since methionine-derived methyl groups are required for phospholipid and lipoprotein synthesis ^25^ and methionine availability also impacts cholesterol synthesis rate ^26^. We also observed subpopulation-specific gene expression changes, notably increase of the epithelial cell cadherin CDH1 in Cluster 0 but not Cluster 1 in revertant cells (Fig. 3f), further indicating that cell differentiation state differs between these cell populations. We did not observe any marked expression changes in genes involved in methionine metabolism. Given the requirement for high B_12_ levels for reversion to methionine independence, we also specifically analyzed a set of genes related to B_12_ transport and metabolism. We did not observe concerted expression changes in these pathways, but we did notice changes in B_12_ transporters ABCC1, ABCD4 as well as the B_12_-metabolizing enzyme MMAB (Fig 3f, Suppl Fig 3d), which interestingly has also recently been implicated in regulation of cholesterol homeostasis ^27^.

### Metabolic flux analysis of methionine-dependent cells

We next sought to understand how methionine metabolism differs between methionine-dependent and independent cells, and how such cells respond to methionine substitution. To allow comparisons within an isogenic system, we here used the BJ-TERT and BJ-RAS cell lines. In met^+^ conditions, intracellular methionine was similar in both cell lines and ∼5-fold higher than medium concentrations (Fig 4a), consistent with methionine uptake occurs via a concentrating transporter, while intracellular homocysteine was undetectable (Fig 4b). When cells were subjected to met^−^hcys^+^ medium, these concentrations were drastically altered: methionine dropped 100-fold to low micromolar levels, while intracellular homocysteine increased to medium levels (Fig 4ab). Interestingly, this sharp decrease in methionine content in met^−^hcys^+^ conditions resulted in no more than 5-fold decrease in the central methyl donor S-adenosylmethionine (SAM) levels (Fig 4c), indicating that both cell types strive to maintain sufficient SAM levels. Simultaneously, SAH was increased in met^−^hcys^+^ cultures (Fig 4d), likely due to reversal of the AHCY reaction (Fig. 1a). As a result, the SAM:SAH ratio, which is considered an indicator of methylation potential ^28^, dropped from > 30 in met^+^ conditions to < 5 in met^−^ hcys^+^ (Fig. 4e).

**Figure 4.**
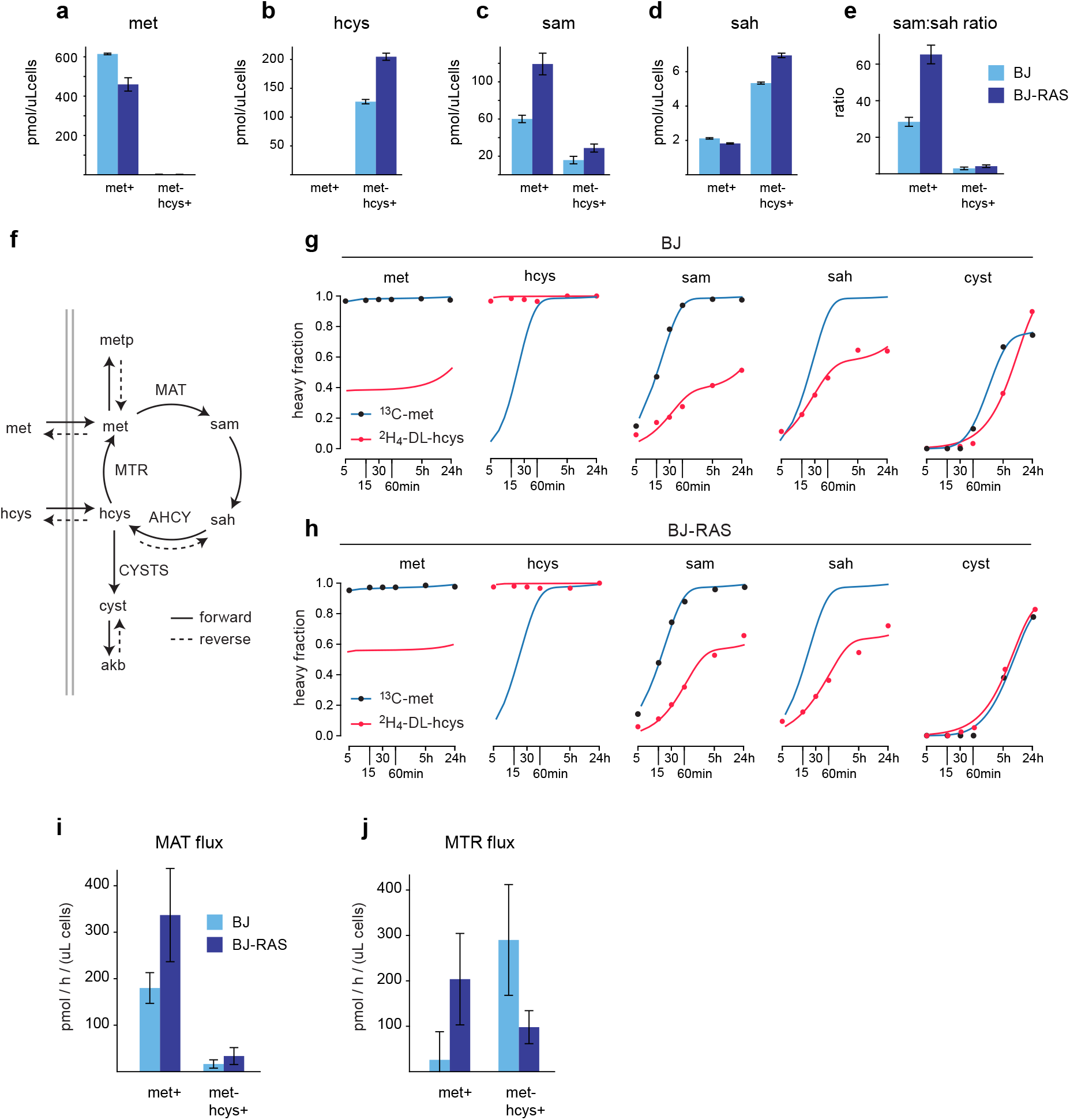
Response of methionine-dependent and -independent cells to homocysteine substitution. **a-d**, Intracellular concentrations of methionine (met), homocysteine (hcys), S-adenosylmethionine (sam) and S-adenosylhomocysteine (sah) in normal fibroblasts (BJ) and isogenic HRAS^V12^ transformed fibroblasts (BJ-RAS), in methionine-containing (met^+^) medium and methionine-free, homocysteine-containing medium (met^−^hcys+). **e**, Ratio of intracellular sam:sah concentrations. **f**, schematic of network model used for metabolic flux analysis. Double gray lines indicate cell membrane. CYSTS, cystathionine synthase; akb, alpha-ketobutyrate; metp, protein-bound methionine; other abbreviations are as in Fig. 1a. **g–h**, Isotope labeling time-course data for indicated metabolites in BJ (g) and BJ-RAS (h) cells, in U-^13^C-methionine (^13^C-met) medium and methionine-free medium containing ^2^H_4_-DL-homocysteine (^2^H_4_-DL-hcys), at indicated time points. Solid lines indicate model fit to data. Heavy (isotope-labeled) fraction is shown; see Methods for details. **i–j**, Estimated flux through the MAT (i) and MTR (j) reactions in BJ and BJ-RAS cells, in indicated media.

To gain more insight into methionine metabolism in these conditions, we performed time-series isotope tracing experiments with ^13^C_5_-methionine in met^+^ medium or ^2^H_4_-DL-homocysteine in met^−^hcys^+^, and estimated metabolic fluxes using model-based flux analysis (Fig 4f; see Methods). In met^+^ cultures, intracellular methionine was fully ^13^C-labeled already at 5 minutes (Fig 4g), indicating rapid exchange with the medium. Overall, measured and model-fitted isotope labeling dynamics were very similar in the two cell lines (Fig. 4g,h, Supplementary Fig. 4a,b). In particular, SAM half-life was consistently around 15 minutes; yet, MAT flux was higher in BJ-RAS cells (Fig 4i) due to a larger SAM pool in these cells, in line with previous reports of increased MAT flux in transformed cells ^29^. When subjected to met^−^hcys^+^ medium, SAM isotope labeling (Fig. 4g,h) and MAT flux (Fig. 4h) was markedly reduced in both cell types, consistent with the low SAM:SAH ratio. Exchange flux through the reversible AHCY reaction also increased markedly, evidenced by rapid labeling of SAH from homocysteine. Flux from homocysteine into cystathionine was very small in all cases (Supplementary Fig. 3c,d), indicating that the “transsulfuration” pathway is not quantitatively important in these conditions.

Unfortunately, flux through the MTR reaction cannot be directly measured using isotope tracing, since rapid exchange of intracellular methionine with the much larger medium methionine pool means that the isotopic state of intracellular methionine is virtually always the same as that of medium methionine, in effect “erasing” isotopic information on MTR activity. Instead, we estimated MTR flux indirectly from mass balance constraints in the model, exploiting the fact that methionine consumed for methylation and protein synthesis must equal methionine uptake plus synthesis via MTR. In BJ-TERT cells, MTR flux was increased in met^−^hcys^+^ conditions (Fig. 4j), reflecting sustained methionine demand in the absence of methionine uptake. In contrast, methionine-dependent BJ-RAS cells exhibited low MTR flux in met^−^hcys^+^ medium (Fig. 4j). Although these flux estimates are uncertain, this data nevertheless suggests potential differences in MTR between methionine-dependent and independent cells.

### Loss of methionine synthase activity underlies methionine dependence

Given that high B_12_ promotes methionine reversion and that MTR flux appears to be altered in methionine-dependent BJ-RAS cells, we wondered if growth of cancer cells in met^−^hcys^+^ condition could be limited by insufficient MTR activity. To test this hypothesis, we first attempted to shift the MTR reaction towards methionine synthesis by culturing cells in medium with 10-fold higher homocysteine (1 mM). Interestingly, this partially rescued growth of both tumor-derived cancer cells (Fig 5a,b) and transformed fibroblasts (Fig 5c). In this condition, intracellular homocysteine increased to > 1mM (Fig. 5d). Although intracellular methionine was not detectable, intracellular SAM levels increased ∼5-fold (Fig 5e), suggesting that methionine cycle function was partially restored. However, SAH also increased to very high levels (Fig 5f), likely due to backflux through the AHCY enzyme caused by the high homocysteine concentration, and consequently the SAM:SAH ratio was not restored (Fig 5g).

**Figure 5.**
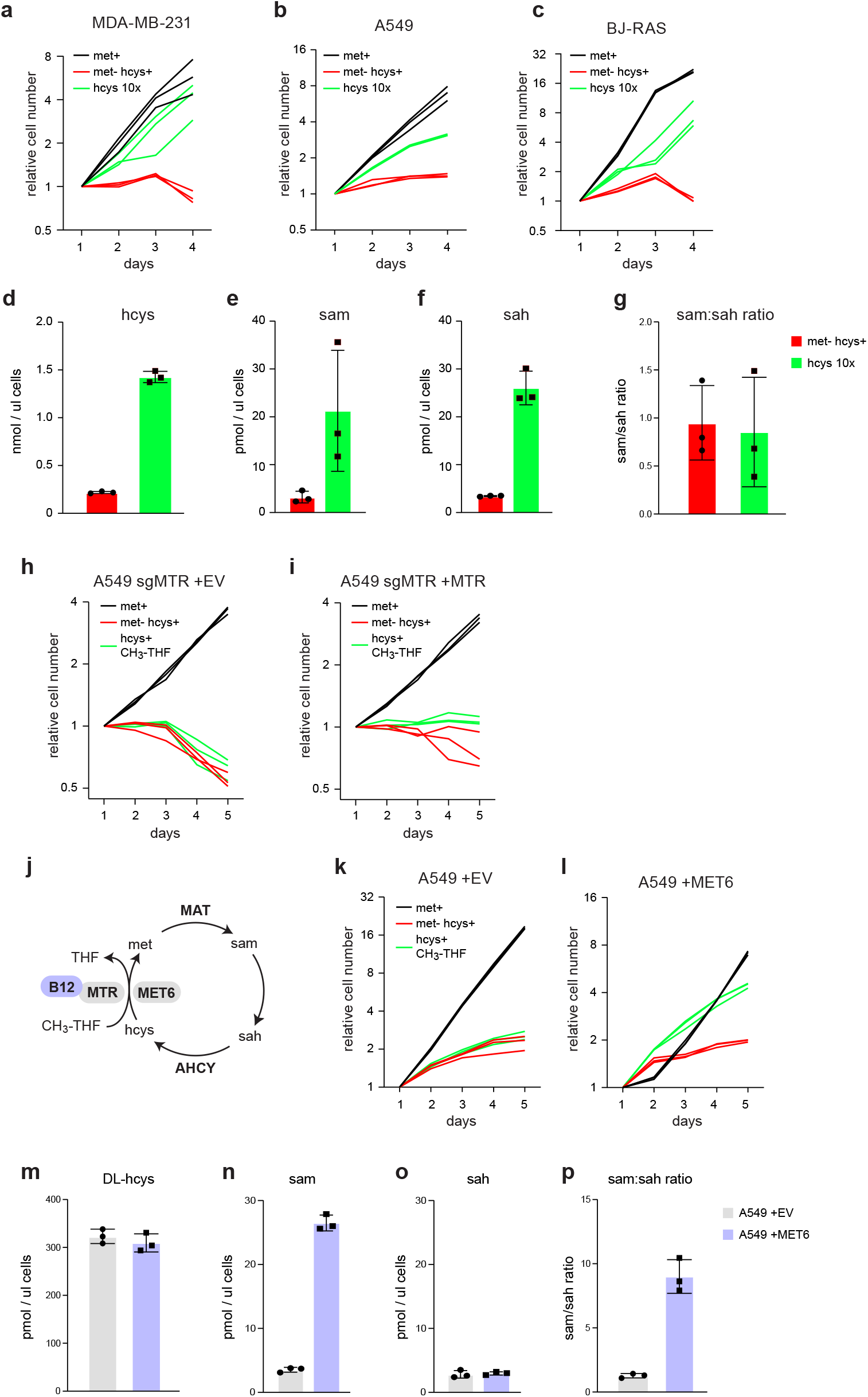
Rescue of methionine dependence in cancer cells by methionine synthase. **a–c**, Growth curves of MDA-MB-231 (a) A549 (b) and BJ-RAS (c) cells in methionine-containing (met^+^) medium, methionine-free medium with 100 uM homocysteine (met^−^hcys^+^), and methionine-free medium with 1 mM homocysteine (hcys 10x). **d–f**, Intracellular concentrations of homocysteine (hcys), S-adenosylmethionine (sam) and S-adenosylhomocysteine (sah) in A549 cells cultured in indicated media. **g**, Ratio of intracellular sam:sah concentrations. **h–i**, Growth curves of A549 cells with CRISPR knockout of endogenous MTR (sgMTR), over-expressing empty vector (+EV; h) or MTR (+MTR; i), in met^+^ or met^−^hcys^+^ media, or met^−^hcys^+^ media supplemented with CH_3_-THF. **j**, Schematic of methionine cycle B_12_-dependent MTR and the B_12_-independent methionine synthase MET6 indicated. **k–l**, Growth curves of A549 cells over-expressing empty vector (+EV; k) or MET6 (+MET6; l), in indicated media. **m–o**, Intracellular concentrations of homocysteine (hcys), S-adenosylmethionine (sam) and S-adenosylhomocysteine (sah) in A549 cells cultured in indicated cell lines in met^−^hcys^+^ medium. **p**, Ratio of intracellular sam:sah concentrations. Cell numbers relative to day 1 from three independent time course experiments are shown (n = 3). Error bars denote standard deviation (n=3).

To test whether methionine dependence might be due to insufficient MTR expression, we turned to a previously established model ^30^ where MTR was overexpressed on a background of CRISPR-Cas9 MTR knockout cells. Overexpression of MTR in this model failed to improve growth of cells in met^−^hcys^+^ conditions, even with provided the MTR substrate 5-methyltetrahydrofolate (CH_3_-THF; Fig. 5h,i), and in the presence of high B_12_ (Supplementary Fig 5a,b). We reasoned that this might be due to failure to increase levels of functional MTR enzyme, which requires insertion and reduction of the B_12_ prosthetic group, a complex process requiring several accessory proteins ^31^. To circumvent this difficulty, we decided to express a B_12_-independent methionine synthase MET6 from *S. cerevisiae* in A549 cells, which should allow cells to synthesize methionine from homocysteine (Fig 5j) in the setting of insufficient B_12_. Remarkably, MET6 expression (Supplementary Fig. 5c) restored robust cell growth in met^−^hcys^+^ medium, provided that CH_3_-THF was added to the medium (Fig 5k,l). Hence, insufficient supply of endogenous CH_3_-THF may contribute to methionine dependence in these cells. Moreover, MET6 expression increased intracellular SAM concentrations while SAH was unaffected (Fig 5n,o), and thereby restored the SAM:SAH ratio (Fig 5p). We conclude that methionine dependence in these cells is due to insufficient methionine synthase activity related to B_12_ deficiency.

## Discussion

Taken together, our results indicate that methionine dependence in cancer cells is due to loss of MTR activity, at least in the cell lines studied. This contrasts with the commonly cited theory that methionine-dependent cells have fully functional methionine synthesis ^13,32^. According to this theory, sufficient methionine is formed by MTR, but this methionine is somehow distinct from exogenous methionine in that it cannot be used by methionine adenosyltransferase (MAT; Fig 1a) to form S-adenosylmethionine (SAM) ^14^. Why this would be the case is unclear, given that both the MTR and MAT enzymes are present in the cytosol ^33^ and that MTR-derived methionine is evidently a substrate for MAT in methionine-independent cells. To our knowledge, the evidence for sufficient methionine synthesis in methionine-dependent cells consists mainly of measurements of enzymatic activity assays in cell lysates ^9,13^, performed using high levels of B_12_ and reaction substrates, which do not reflect MTR flux in living cells. Indeed, fibroblasts with genetic defects in B_12_ metabolism that disable methionine synthesis *in vivo* can appear normal in such assays ^34^. On the other hand, one study of glioma cell lines reported low B_12_ levels and reduced MTR activity in met-dependent cells ^19^, in line with our results. Considering these points, and in the light of our findings, we propose that loss of methionine synthase activity is the most parsimonious explanation for methionine dependence in cancer cells.

Several studies have highlighted the importance of SAM, the central methyl donor in mammalian cells, as a key mediator of methionine dependence. SAM is depleted in cells starved of methionine ^16,29^, and loss of SAM leads to cell cycle arrest via a “checkpoint” machinery ^35^. One study also reported that addition of SAM enabled methionine-dependent MDA-MB-468 cells to grow in met^+^hcys^−^ medium ^16^. Our finding that loss of MTR activity underlies methionine dependence does not contradict these results, but suggests that insufficient SAM is secondary to lack of methionine synthesis. Notable, we observe a marked drop in SAM and in the SAM:SAH ratio in methionine-dependent cells subjected to met^+^hcys^−^ medium, which is reversed by expression of methionine synthase. Others have suggested that transformed cells might have an increased demand for SAM to drive cellular methylation ^29^, which may make them more sensitive to loss of methionine. We did find somewhat higher SAM synthesis rates in RAS-transformed cells, but whether this difference is quantitatively important is not clear.

The underlying cause of low MTR activity in cancer cells remains to be determined. The fact that the B_12_-independent methionine synthase MUT6, but not expression of human MTR, rescues the methionine dependence phenotype strongly suggests that the defect lies in providing a functional B_12_ cofactor to the MTR enzyme. That high B_12_ concentration facilitates adaptation to met^−^hcys^+^ medium, which has apparently been discovered by several laboratories (Suppl Table 1) further strengthens this notion. In addition, one report indicated that MTR isolated from methionine-dependent cells occurs mostly in the apoenzyme form (lacking the B_12_ cofactor), unlike MTR from methionine-independent normal cells ^36^. Importantly, a functional B_12_ deficiency would be expected to be tolerated in cancer cells in met^+^ conditions (including *in vivo*), since the only human enzymes that require B_12_ are MTR and MUT, of which MUT is only necessary for propanoate oxidation and MTR appears to be dispensable for growth in normal nutrient conditions ^30^. It therefore seems plausible that B_12_ deficiency could arise in tumors.

Some important caveats should be mentioned. While our metabolic flux analysis indicates reduced MTR flux in methionine-dependent cells in met^−^hcys^+^ medium, we emphasize this is only an indirect estimate based on mass-balance considerations. Direct measurement of MTR flux in living cells is very difficult since the reaction is not observable with typical isotope-tracing methods due to rapid equilibration of methionine across the cell membrane, a problem that may have obscured MTR defects in methionine dependence until now. Our flux analysis also did not consider consumption of SAM by other processes than methylation, such as polyamine synthesis, and may therefore overestimate the cellular methylation rate. Importantly, while our data shows that overexpression of methionine synthase rescues growth of methionine-dependent cells, this requires that CH_3_-THF is provided to cells. Therefore, in addition to having insufficient MTR activity, these cells appear unable to generate enough CH_3_-THF to support methionine synthesis. It is currently not clear how these two aspects of methionine deficiency are related, and this is an important avenue for future research.

The observation that a variety of cancer cell types are strongly dependent on methionine, and cannot easily adapt to grow without it, naturally suggests that dietary methionine restriction could be an effective strategy for cancer therapy. Our observation from scRNA-seq data that homocysteine selection favors a less aggressive phenotype also supports this idea. While this approach has been effective in mouse models ^3–8^, methionine restriction is not well tolerated in humans, and has yet to be successful in clinical trials. Our findings may lead to new ways of refining this approach. For example, if methionine-dependent cancers are generally B_12_-deficient, then a test for B_12_ status might identify individual patients that could benefit from methionine deprivation. Also, dietary interventions could perhaps be modified to maintain homocysteine levels in a range that supports methionine synthesis in normal tissues, to mitigate adverse effects. In any case, by establishing a biochemical basis for methionine dependence, we hope that our results will help exploit this metabolic phenomenon for cancer therapeutics.

## Methods

### Cell culture

BJ-TERT, BJ-SV40, BJ-RAS, MDA-MB-231 (HTB-26, ATCC), HMEC (CC-2551, Lonza), and A549 (CCL-185, ATCC) were cultured under specific conditions. BJ-TERT, BJ-SV40, and BJ-RAS were grown in RPMI 1640 medium (catalog: 61870010, ThermoFisher Scientific), supplemented with 5% heat-inactivated fetal bovine serum (FBS) (catalog: 16140071, Gibco) and 1% penicillin-streptomycin (catalog: 15140122, Gibco). MDA-MB-231 and A549 were maintained in RPMI 1640 medium with 10% FBS and 1% penicillin-streptomycin. HMEC cells were cultured in MCDB170 medium (catalog: M2162-06, USBiological Life Sciences), supplemented with Mammary Epithelial Growth Supplement (MEGS) (S0155, ThermoFisher), glucose (8.00E-03M, 1181302, Sigma), and glutamine (2.00E-03M, 49419, Sigma).

### Custom medium synthesis

Custom RPMI 1640 and MCDB170 media were prepared in-house and by omitting methionine and supplementing with 30uM, 100uM or 1mM homocysteine (69453, Sigma-Aldrich) for MCDB170 (Met-Hcys+), RPMI 1640 (Met-Hcys+), 10x homocysteine RPMI (Met-10x Hcys+) respectively as outlined in Supplementary Table 4. Custom RPMI 1640 was supplemented with heat-inactivated fetal bovine serum (FBS) (catalog: 16140071, Gibco) dialyzed in SnakeSkin 3,500 molecular-weight cut-off dialysis tubing (88244, Thermo Fisher) and 1% penicillin–streptomycin (PeSt; Gibco,15140148). MCDB170 was supplemented with Mammary Epithelial Growth Supplement (MEGS) (S0155, ThermoFisher), glucose (8.00E-03M, 1181302, Sigma), and glutamine (2.00E-03M, 49419, Sigma). For hcys+B12 or hcys+B12+5mTHF medium, 1.5uM of vitamin B12 (cyanocobalamin, V6629, Sigma-Aldrich) and/or 4.4uM 5 methyl tetrahydrofolate was added (M0132, Sigma-Aldrich). For tracing the same custom medium was supplemented with 100uM U-^13^C_5_-L-methionine (Cambridge Isotope Laboratories, CLM-893-H) at 99% atom purity or 200uM with 3,3,4,4,-^2^H_4_-DL-homocystine (Cambridge Isotope Laboratories, DLM-8259)

### MET6 overexpression experiments

For overexpression of the B_12_-independent methionine synthase from yeast (MET6) ^37^, the coding sequence (CDS) from refseq sequence no. NM_001178982.3, was codon-optimized for mammalian expression and synthesized to ensure efficient expression in human cells. The length of the codon-optimized MET6 CDS is 2304 base pairs (bp). Lentiviral vector construction, production, and transduction were conducted by Cyagen Biosciences in Santa Clara, California, USA. Two groups of lentiviral vectors were constructed: the experimental group consisted of LV-EF1A>S. cerevisiae MET6 CDS [NM_001178982.3], which included a Kozak sequence and the EF1A promoter along with mPGK>Puro as the selection marker, while the control group used LV-mPGK>Puro with puromycin as the selection marker. The expression of MET6 was quantified using RT-qPCR with primers specific to the codon-optimized MET6 transcript.

### Methionine reversion

MDA-MB-231, MCF7, BJ-RAS, and A549 were grown to approximately 60-70% confluency in 6 well plates. At this stage, the methionine-containing medium was replaced with methionine-free medium supplemented with homocysteine. <u>M</u>edium was replaced daily to remove dead cells during the first week and every 2 days during the second week. During the third week, medium was replaced every three days to maintain colony stability and minimize disturbance to the growing cells. Cell confluency was monitored throughout the experiment using IncuCyte S3 Live-Cell Imaging System (Sartorius AG, Göttingen, Germany). At the end of the reversion experiment cells were collected and counted using Sceptre 3 (Millipore).

### Proliferation assays

Cells were seeded into 24- or 48-well plates (Sarstedt) at the following densities: MDA-MB-231 (2 × 104 cells/cm^2^), A549 (1 × 104 cells/cm^2^), BJ-TERT (1 × 104 cells/cm^2^), BJ-RAS (5 × 10^3^ cells/cm^2^), HMEC (2.5 × 10^3^cells/cm^3^), GB11 (1 × 104 cells/cm^2^), MCF7 (2 × 104 cells/cm^2^), and HCT116 (2 × 104 cells/cm^2^). For methionine dependence characterization, cells were cultured in methionine-containing (Met), homocysteine-containing (Hcys), or methionine + homocysteine (Met + Hcys) medium (Fig. 1, Fig. S1). After reversion, cells were reseeded in Met, Hcys, or Hcys + B12 medium to assess growth post-reversion. For A549 variant cell lines (A549-EV, A549-MTR, and A549-MET6), cells were seeded at 1 × 104 cells/cm^2^.

For cell counting, plates were imaged every 24 hours, and cell counts were obtained using a deep learning-based segmentation classifier trained to recognize the cell types involved. The classifier software is available at https://github.com/Nilsson-Lab-KI/unet-cell-counting. Cell doubling times were calculated from initial and final number assuming exponential growth.

### Single-cell RNA sequencing

Cells were collected and counted using a hand-held Coulter counter (Scepter 3, Merck Millipore). Cells were diluted to a target concentration of approximately [1000 cells/µL] for optimal loading onto the 10x Genomics Chromium Controller. Single-cell suspensions were processed using the 10x Genomics Chromium Next GEM Single Cell 3’ Kit v3.1 (CG000315 REV E) according to the manufacturer’s protocol. Cells were loaded onto the Chromium Controller to generate Gel Bead-In Emulsions (GEMs) containing individual cells, barcoded primers, and enzymes necessary for reverse transcription (PN-1000123, 10x Genomics). GEM generation and barcoding efficiency were monitored during the process to ensure high-quality libraries. GEMs were lysed to release cellular RNA, followed by reverse transcription and amplification to generate full-length cDNA (PN-1000190, 10x Genomics). cDNA libraries were fragmented, end-repaired, A-tailed, and ligated to sequencing adapters. Library quality and concentration were assessed using the Agilent Bioanalyzer 2100 (catalog) and Qubit dsDNA HS Assay Kit (catalog), respectively. Libraries were sequenced-ready after achieving a target fragment size of approximately [∼500 bp]. Libraries were sequenced on an Illumina [NovaSeq 6000/NextSeq] platform using paired-end reads with a read length of 150 bp.

### Isotope tracing

For tracing with ^13^C_5_-methionine, medium contained 100uM U-^13^C_5_-L-methionine (Cambridge Isotope Laboratories, CLM-893-H) at 99% atom purity. For homocysteine tracing, medium contained 200uM with 3,3,4,4,-^2^H_4_-DL-homocystine (Cambridge Isotope Laboratories, DLM-8259), corresponding to 100uM L-homocysteine, and no methionine. The tracer purity as reported by the vendor was 99.6%. In each case, cells were precultured for 4 hours in an unlabeled medium of the same composition but with unlabeled methionine and homocysteine, respectively, and switched to the corresponding isotope-labeled medium at time 0. Cell extracts were harvested at 5, 15, 30 min, and 1, 5 and 24 h. Medium samples (supernatants) were taken at the final 24 h timepoint. In addition, medium incubated without cells for 24 h was used as control for uptake/release measurements.

### Concentration and uptake/release measurements

Concentrations of were measured in isotope-labeled culture medium and in cell extracts using isotope dilution with internal unlabeled standards, as previously described ^38^. Briefly, an amount *n*_*std*_ of a pure standard with mass isotopomer (MI) fraction *x*^*std*^ was added to an unknown amount *n* of the corresponding metabolite in cell extracts with MI fraction *x*, and the MI fraction *x*^*mix*^ of the resulting mixture was measured. It then holds that

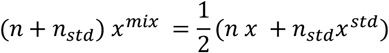

from which we can solve for the unknown *n*. To convert cell extract concentrations to intracellular concentrations, the total cytosol volume of extracted cells was estimated from cell diameters measured using a Coulter counter (Scepter 3.0, Merck Millipore). Metabolite release rates were computed by subtracting concentrations in medium incubated without cells from concentrations in spent medium.

### Metabolic flux analysis

For metabolic flux analysis, we used a simplified model where each metabolite is considered as a mixture of “light” (unlabeled) or “heavy” (labeled) forms, and the methionine metabolic network is represented as a single compartmental system. The system is then fully described by a differential equation system with a single variable for each metabolite, representing the heavy fraction. In general, an isotope labeling system can be expressed in this compartmental form if each metabolite in the system has only one labeled isotopomer (after correction for natural abundance and isotopomer impurities). For the ^2^H_4_-homocysteine tracing experiments, this holds true since the ^2^H_4_-labeled moiety is not altered by any of the reactions in the model. For the U-^13^C_5_-methionine tracing experiments, the assumption would be violated if ^13^C_4_-methionine produced by MTR was substantial; however, since measured ^13^C_4_ mass isotopomer of methionine was negligible, the forward rate of ^13^C_4_-methionine production by MTR is negligible compared to the influx of methionine, and the methionine pool can be modeled as a mixture of ^13^C_5_ and ^13^C_0_ isotopomers.

To estimate the “light” and “heavy” isotopomer fraction for a metabolite, we fit a linear mixture model

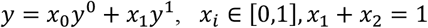

where *y*^0^ is the natural abundance MID and *y*^1^ is the expected fully labeled MID at a given atom purity (both binomial distributions). The above equation was fit to observed data by the standard least-squares method, and the heavy fraction *x*_1_ was used for metabolic flux analysis. Atom purity was > 99% in all cases.

Flux estimation was done by fitting the differential equation model to measured heavy fraction time-series, intracellular concentrations and uptake/release fluxes for methionine and homocysteine, plus protein synthesis rate estimated from cell counts and cell size measurements, using the Levenberg-Marquardt method as implemented in the lmfit python package. Goodness-of-fit was evaluated using the chi-square test: for the ^13^C-methionine model, the chi-square values were 13.6 and 17.4 for BJ-TERT and BJ-RAS, respectively with acceptance region (12.0, 21.0); for ^2^H_4_-homocysteine model, the chi-square values were 45.5 and 70.1 for BJ-TERT and BJ-RAS, respectively with acceptance region (18.0, 28.8). Confidence intervals were obtained by linear approximation around the optimal flux values.

A complete specification of the metabolic flux analysis model, all measurement data used, and python code for reproducing the flux analysis is available at https://github.com/Nilsson-Lab-KI/met-hcys-flux

## Acknowledgements

The authors would like to acknowledge Antonio Checa and Peri Noori for support with mass spectrometry analysis and single-cell RNA-sequencing. This work was supported by grants from the Swedish Research Council (2020-01631) and Cancerfonden (20 0974 PjF) to M.E. and R.N.

## Figure legends

**Supplementary Figure 1. Methionine dependence in tumor-derived cancer cells. a–c**, Growth curves for colon cancer cells (HCT116), breast cancer cells (MCF7), lung cancer cells (A549) and glioblastoma cells (GB11) in methionine-containing (met^+^) medium and methionine-free, homocysteine-containing medium (met^−^hcys^+^). **d–f**, Growth curves for MCF7 cells, breast cancer cells (MDA-MB-231) and BJ cells transformed with the SV40 Large-T antigen and oncogenic HRAS^V12^ (BJ-RAS), in met^+^ medium or mediun containing both methionine and homocysteine (met^−^hcys^+^). Cell numbers relative to day 1 from three independent time course experiments are shown (n = 3).

**Supplementary Figure 2. Reversion of methionine dependence in RAS-transformed cells. a**, Long-term growth curves of BJ cells transformed with the SV40 Large-T antigen and oncogenic HRAS^V12^ (BJ-RAS) in homocysteine-containing medium with either 0.003 µM (met^−^hcys^+^) or 1.5 µM vitamin B_12_ (met^−^hcys^+^ B_12_). Cell numbers in two independent cultures are shown (n=2). **b**, Growth curves of revertant cells BJ-RAS(R) obtained from long term cultures, in methionine-containing (met^+^) medium and methionine-free, homocysteine-containing medium (met^−^hcys+). Cell numbers relative to day 1 from three independent cultures are shown (n = 3).

**Supplementary Figure 3. Gene expression patterns in parental and revertant cells. a**, Non-redundant gene ontology (GO) pathways enriched in cluster 2, visualized as UMap projection of semantic similarity. Size of circles represent pathway over-representation score. **b**, S-plot of metastatic signature versus ratio of gene expression in cluster 0 over cluster 1. **c**, p-values for expression signature match against human 1,355 cell types in cluster 0 and cluster 1. Selected cell types are highlighted. **d**, Ratio of gene expression levels in D21 over D0 cells for selected genes involved in B_12_ transport and metabolism.

**Supplementary Figure 4. Metabolic flux analysis in methionine-dependent and -independent cells. a-b**, Model predictions for time-course isotope labeling for non-measured metabolites alpha-ketoburytate (akb) and protein-bound methionine (metp) in BJ (a) and BJ-RAS (b) cells, in U-^13^C-methionine (^13^C-met) medium and methionine-free medium containing ^2^H_4_-DL-homocysteine (^2^H_4_-DL-hcys), at indicated time points. **c**, Intracellular concentrations of cystathionine (cyst) in BJ and BJ-RAS cells, in methionine-containing (met^+^) medium and methionine-free, homocysteine-containing medium (met^−^hcys+). **d**, Estimated flux through the CYSTS reaction in BJ and BJ-RAS cells, in indicated media.

**Supplementary Figure 5. Cancer cell growth rescue and MET6 overexpression. a–b**, Growth curves of A549 cells over-expressing empty vector (+EV; k) or MET6 (+MET6; l), in methionine-containing (met^+^) medium; methionine-free, homocysteine-containing medium (met^−^hcys+); and in met^−^hcys+ medium supplemented with CH_3_-THF and vitamin B_12_. **c**, qPCR of MET6 expression in A549 +EV and A549 +MET6 cels. Data is presented as 2^-ΔΔCt^ from 3 replicates.

