## Supplementary figures and images for "Methionine dependence in cancer cells due to lack of B_12_-dependent methionine synthase activity"

### Supplementary Figure 1

##### Supplementary Figure 1

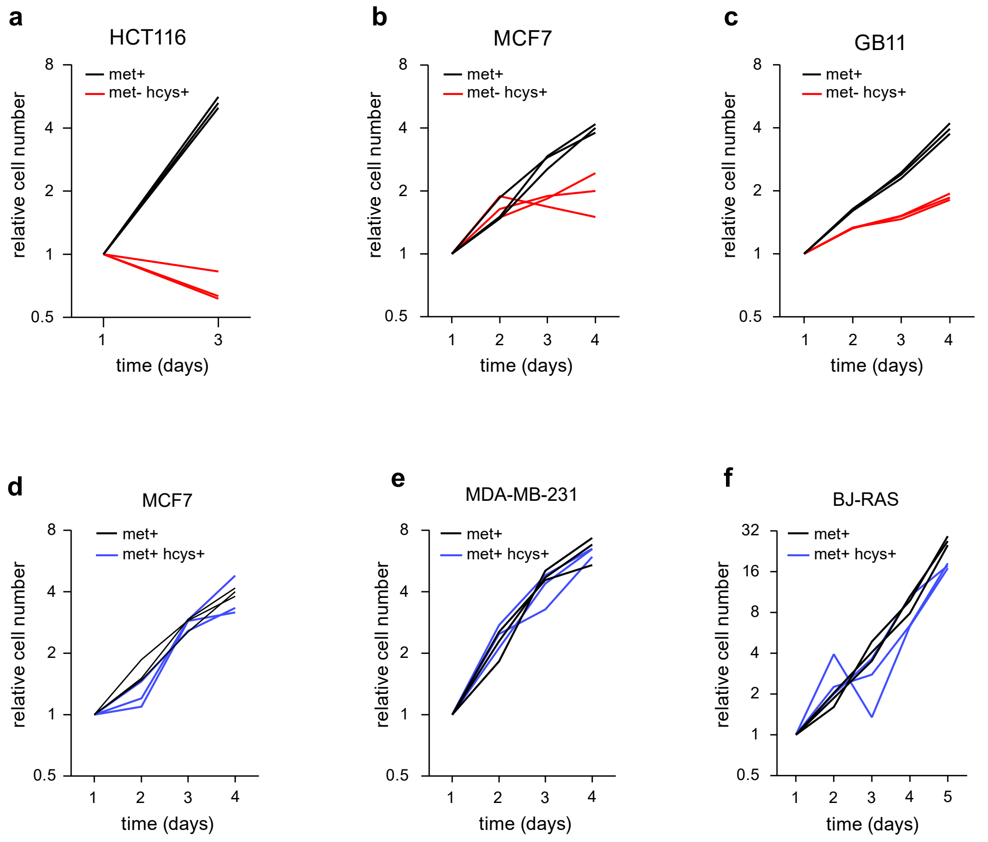

### Supplementary Figure 2

**Supplementary Figure 2**

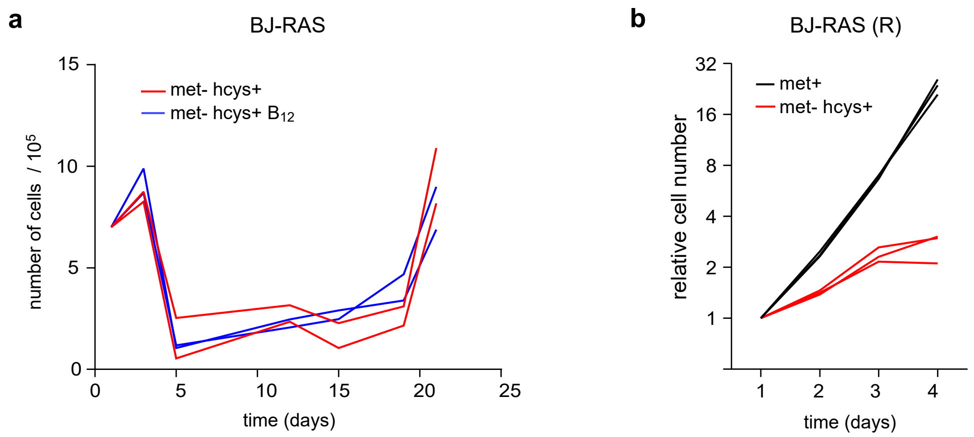

### Supplementary Figure 3

#### Supplementary Figure 3

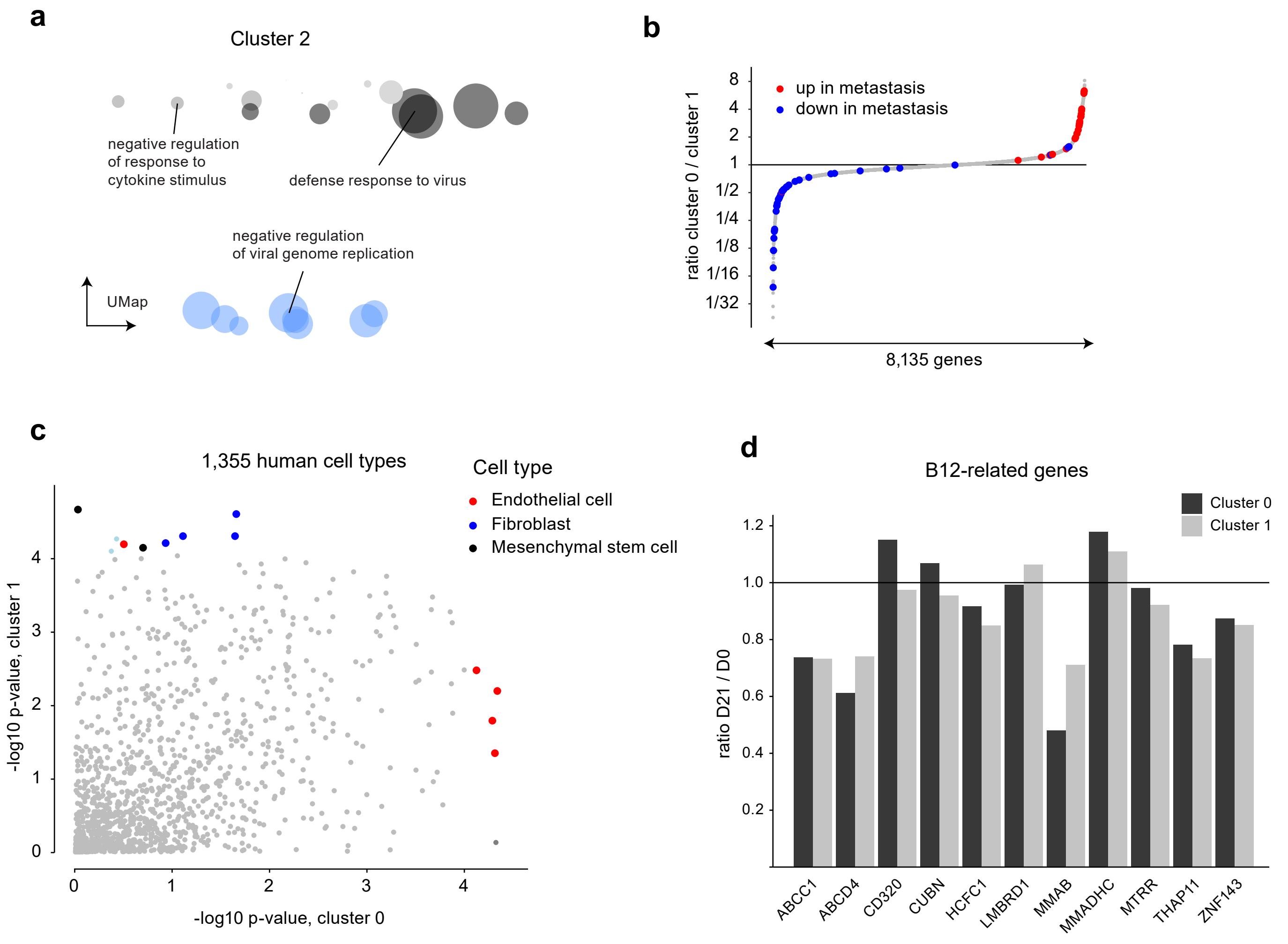

### Supplementary Figure 4

#### Supplementary Figure 4

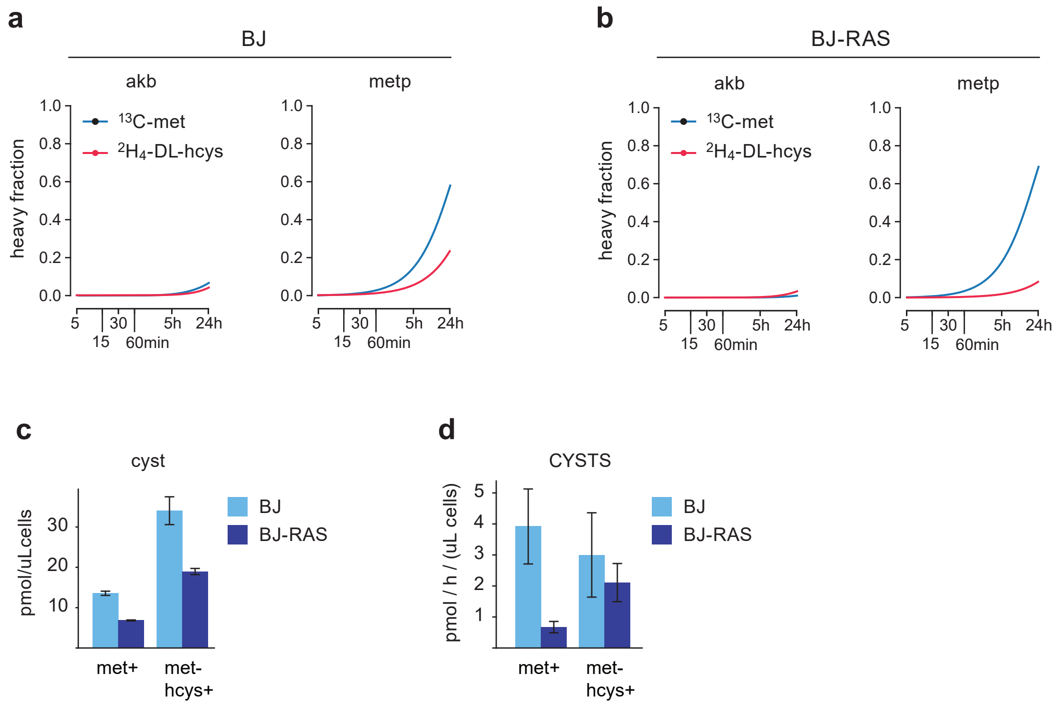

### Supplementary Figure 5

#### Supplementary Figure 5

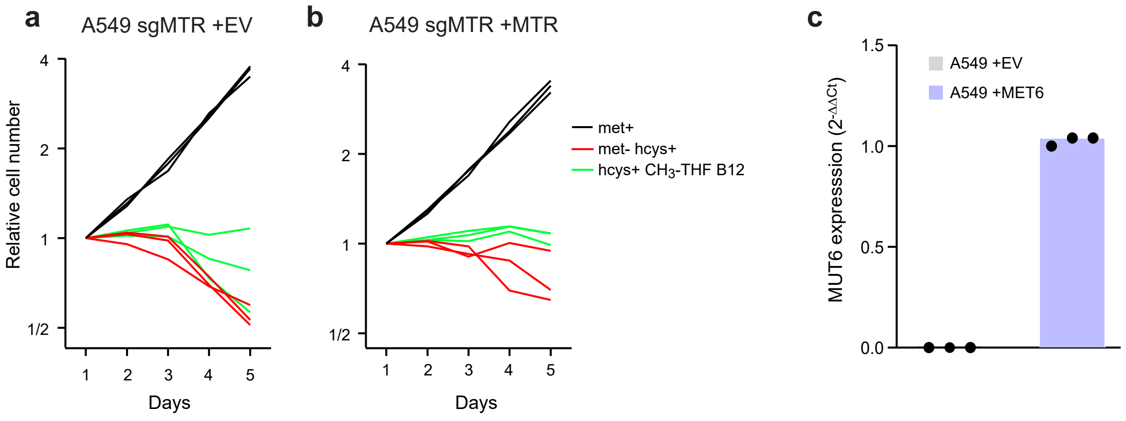
